# The Effect of P2X7 Antagonism on Subcortical Spread of Optogenetically-Triggered Cortical Spreading Depression and Neuroinflammation

**DOI:** 10.1101/2022.09.26.509535

**Authors:** Burak Uzay, Buket Donmez-Demir, Sinem Yilmaz Ozcan, Emine Eren Kocak, Muge Yemisci, Yasemin Gursoy Ozdemir, Turgay Dalkara, Hulya Karatas

## Abstract

Migraine is a neurological disorder characterized by episodes of severe headache. Cortical spreading depression (CSD), the electrophysiological equivalent of migraine aura, results in opening of pannexin-1 megachannels that release ATP and triggers parenchymal neuroinflammatory signaling cascade in the cortex. Migraine symptoms suggesting subcortical dysfunction bring subcortical spread of CSD under the light. Here, we investigated the role of purinergic P2X7 receptors on the subcortical spread of CSD and its consequent neuroinflammation using a potent and selective P2X7 antagonist, JNJ-47965567. P2X7 antagonism had no effect on the CSD threshold and characteristics but increased the latency to hypothalamic voltage deflection following CSD showing that ATP acts as a mediator in the subcortical spread. P2X7 antagonism also prevented hypothalamic neuronal activation following CSD, revealed by bilateral decrease in hypothalamic c-fos positive neuron count. P2X7 antagonism further stopped the CSD-induced neuroinflammation revealed by decreased nuclear translocation of NF-kappa B-p65 in astrocytes and decreased HMGB1 release. Following CSD we observed an increase in neuronal cytoplasmic P2X7R signal in cortex and subcortical structures (thalamus, hypothalamus, striatum, hippocampus) concordant with the neuroinflammation which is also prevented by P2X7R antagonism. In conclusion, our data suggest that P2X7R plays an imperative role in CSD-induced neuroinflammation, subcortical spread of CSD and CSD-induced hypothalamic neuronal activation hence can be a potential target in migraine treatment.

## Introduction

Migraine is a primary episodic headache disorder that affects 10-20% of the population and has a big negative impact on the daily functioning of patients^1^. The pathophysiology of migraine is yet to be fully elucidated but the *cortical spreading depression* (CSD) is known to have a vital role in the pathophysiology of migraine. CSD is the slowly spreading depolarizing electrical activity on the cortex and is the electrophysiological correlate of the migraine aura^2–4^. Previous research has showed that CSD results in opening of the neuronal pannexin-1 (Panx-1) channels, release of proinflammatory cytokines and alarmins (such as HMGB1 and IL1β) and activation of the inflammatory cascade by translocation of p65 subunit of NFκB to the nucleus in the parenchymal astrocytes^5^. This parenchymal inflammation activates meningeal nociceptors via glia limitans and the trigeminovascular system causing headache^5^. Panx-1 channels are known to be closely associated with purinergic P2X7 receptors that are activated by extracellular ATP and that can further activate the inflammatory cascade, rendering them a target worth investigating in the spread of the CSD and its consequent neuroinflammation^6^. A study showed that genetic loss of P2X7 as part of the P2X7/PANX1pore suppresses spreading depolarization, its inflammatory effects and the trigeminovascular activation in wild-type rats and mice^7^.

CSD spreads to subcortical structures, although it has been almost exclusively investigated as a cortical phenomenon^8–10^. A H_2_^15^O-PET study showed that hypothalamus, along with some other subcortical structures (PAG, Putamen, Caudate nucleus) get activated during a migraine attack^11^. This activation is thought to cause some migraine symptoms suggesting subcortical dysfunction including nausea, sensations of cold/hot, yawning^12^. It is not known whether the subcortical spread of CSD is the reason of these symptoms and whether purinergic P2X7 receptors have a role in this spread.

The conventional CSD induction methods are invasive (electrical stimulation, topical KCl application, pin-prick) and are preceded by a craniotomy^13^. These methods can depolarize neurons but damage vascular structures and astrocytes, thus have non-specific effects. CSD can be triggered non-invasively using optogenetic methods without disrupting the integrity of the skull eliminating the non-specific effects of invasive methods. Therefore, optogenetic induction of CSD is highly useful in investigating CSD-induced neuroinflammation^14^. In this study, to investigate the effects of P2X7 antagonism on cortical and subcortical parenchymal neuroinflammation. we induced CSD using optogenetics, and to investigate the the role of P2X7 receptors on the subcortical spread of CSD, we primarily used the pin-prick method as obtaining subcortical electrophysiology recordings is already invasive. To block P2X7 receptors we used a potent, selective and brain-blood barrier (BBB) permeable P2X7 antagonist, JNJ-47965567^15^. Upon P2X7R antagonist administration, we observed an increase in the latency to subcortical voltage deflection following CSD, and a decrease in hypothalamic c-fos positive neuron count underscoring the role of P2X7 receptors in mediating the subcortical spread of CSD. Moreover, P2X7R antagonism prevented the CSD-induced neuroinflammation, suggesting its detrimental role on migraine pathophysiology and its potential as a novel target in migraine treatment.

## Results

### 1) P2X7 antagonism doesn’t change optogenetic CSD threshold and CSD characteristics

Using the optogenetically-triggered CSD threshold protocol used in (14) (Figure 1b) the average CSD threshold was found to be 17.9 mJ (±2.3 S.E.M; n=15). The minimum energy that could induce CSD was 6 mJ and the maximum was 28 mJ. CSD threshold or amplitude didn’t differ between homozygous (15.7 mJ ± 3 S.E.M; n=9) and heterozygous (21.4 mJ ±3.9 S.E.M; n=7) mice (p=0.31) (Figure 1c-f). Neither P2X7 antagonist (n=7) nor the vehicle administration (n=6) (30% cyclodextrin-sulfobutyl ether sodium salt) had an effect on optogenetically induced CSD threshold or amplitude (p>0.05) (Figure 1g-j).

**Figure 1.**
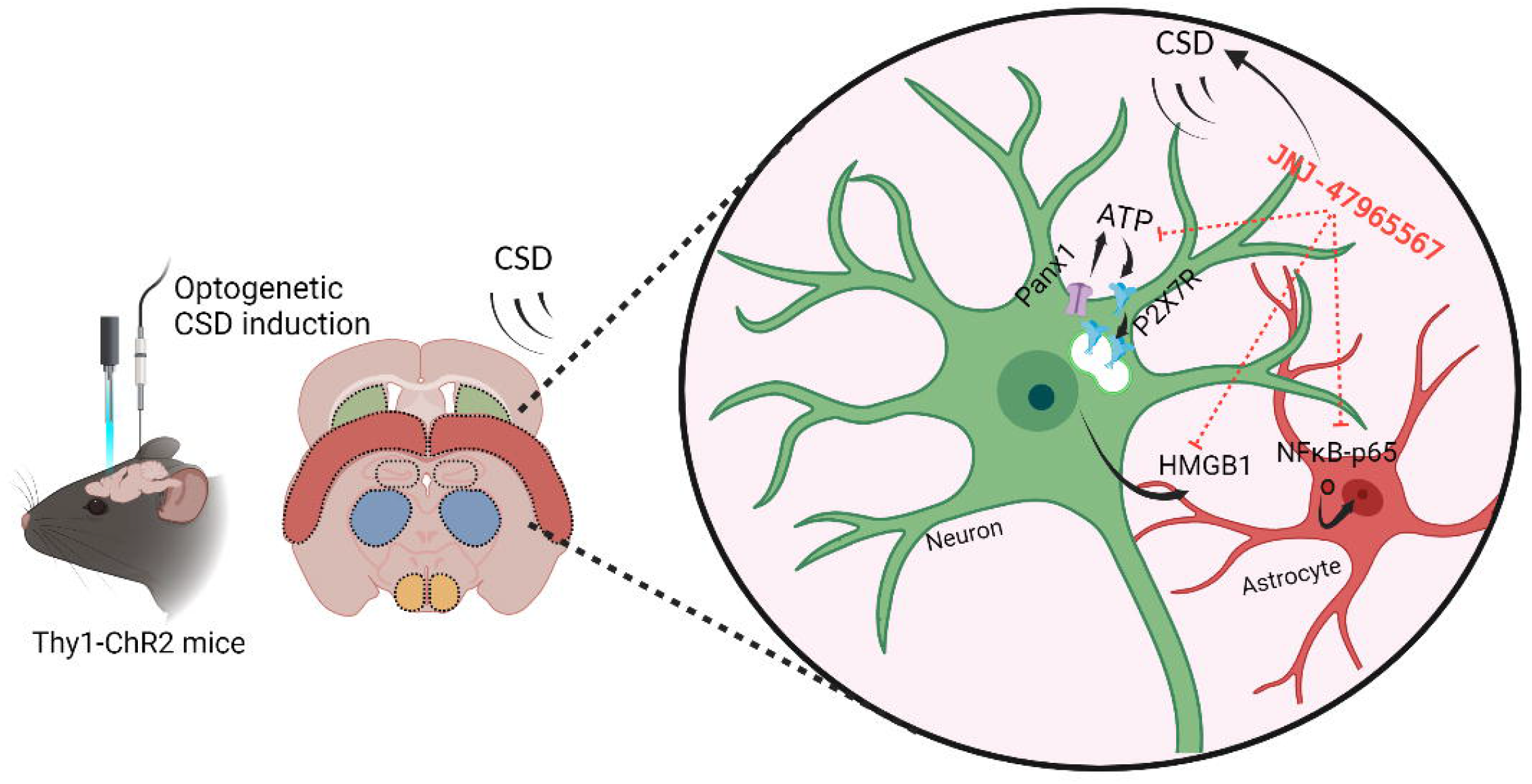
P2X7 antagonism doesn’t change optogenetic CSD threshold and CSD characteristics. **a.** Schematic representation of the experimental protocol **b.** Schematic representation of optogenetic CSD threshold protocol **c.** Representative traces of optogenetically triggered CSD in homozygous(ChR2+/+) and heterozygous(ChR2+/-) mice **d.** Optogenetic CSD Threshold (mJ) in ChR2+/+ (n=9) and ChR2+/-mice (n=7) **e.** CSD amplitude in ChR2+/+ and ChR2+/-mice **f.** Normalized cumulative change in voltage during CSD in ChR2+/+ and ChR2+/-mice **g.** Representative traces of optogenetically triggered CSD in control group and upon P2X7 antagonist or vehicle administration **h.** Optogenetic CSD threshold (mJ) in control group (n=14) and upon P2X7 antagonist (n=7) or vehicle administration (n=6) **i.** CSD amplitude in control group and upon P2X7 antagonist or vehicle administration **j.** Normalized cumulative change in voltage during CSD in control group and upon P2X7 antagonist or vehicle administration. *CSD: Cortical Spreading Depression, ChR2: channelrhodopsin (ns: p>0.05, *:p<0.05,*:p<0.01, *:p<0.001)*

### 2) Subcortical electrophysiological recordings show voltage deflections following CSD

We first performed subcortical electrophysiological recordings from different subcortical structures (striatum, hippocampus, thalamus and hypothalamus) simultaneously with cortical recordings. Upon CSD induction with pin-prick method, a voltage deflection occurred in all investigated subcortical structures (striatum, hippocampus, thalamus and hypothalamus) with varying latencies to CSD (Figure 2a-b).

**Figure 2.**
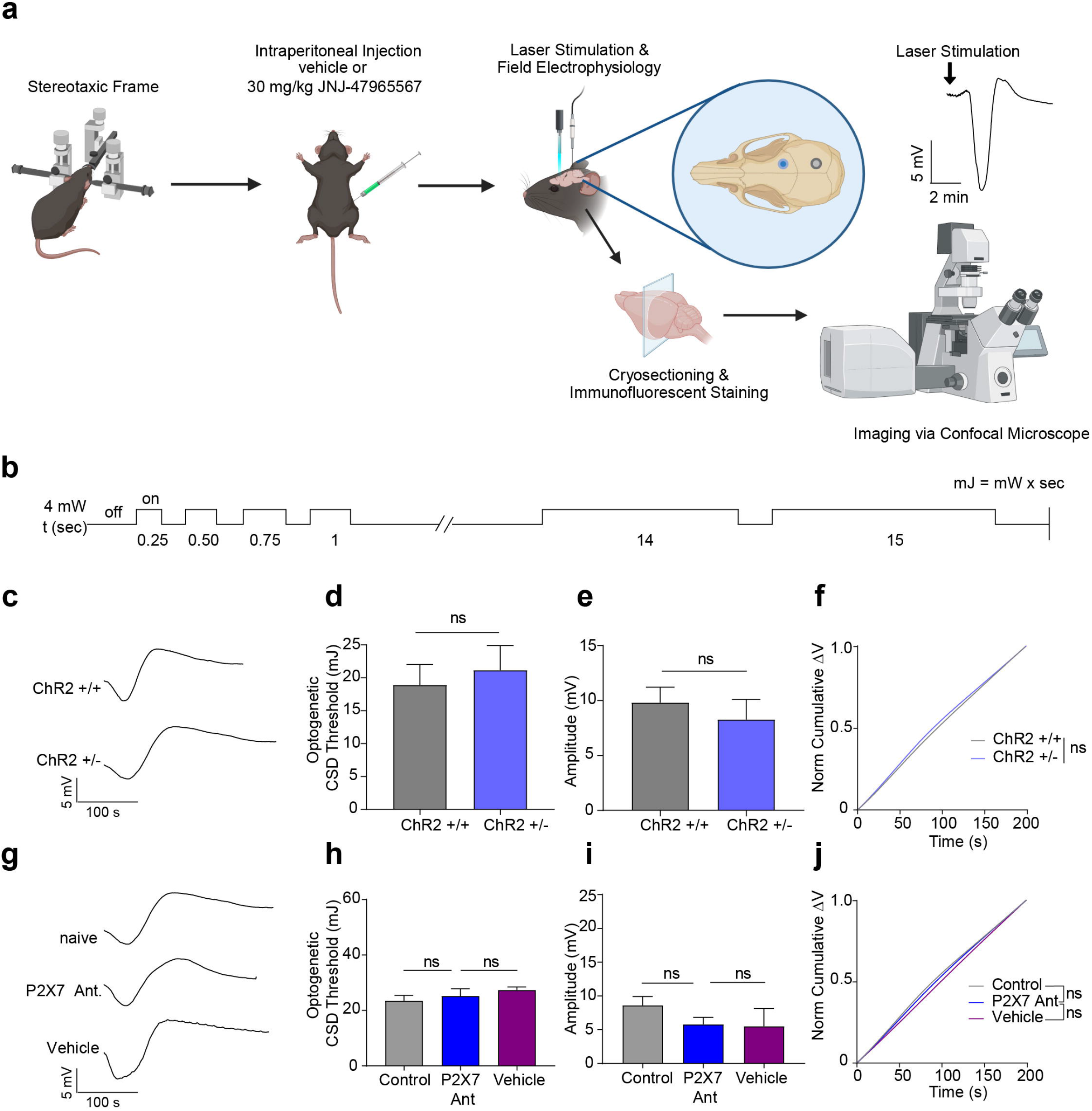
Hypothalamic voltage deflection following CSD happens under P2X7 antagonism with an increase in latency to CSD. **a.** Schematic representation of cortical and subcortical electrophysiology. **b.** Representative traces of cortical(Cx) and subcortical recordings from the, and striatum(Str), hippocampus(Hc), thalamus(Th) and hypothalamus(Hypoth) **c.** Schematic representation of cortical and hypothalamic electrophysiology experimental protocol (green dot: insertion point of tungsten electrode for hypothalamic recordings, pink dot: pin-prick site, grey dot: surface electrode) **d.** Representative traces of cortical and hypothalamic recordings in control group or upon P2X7 antagonist administration. **e.** Hypothalamic voltage deflection latency to CSD in control group and upon P2X7 administration (n=4/group). **f.** Representative images of hypothalamic c-fos immunofluorescent staining in naïve mice, after CSD induction and upon P2X7 antagonist administration. *scale bar:25 μm* **g.** c-fos positive cell numbers in hypothalamus of naïve mice, after CSD induction and upon P2X7 antagonist administration (n=3/group). *CSD: Cortical Spreading Depression (ns: p>0.05, *:p<0.05, *:p<0.01, *:p<0.001)*

### 3) P2X7 antagonism increases the latency of hypothalamic voltage deflection following CSD

Hypothalamic electrophysiological recordings showed a voltage deflection followed by CSD with an average amplitude of 1.05 mV (± 0.18 S.E.M; n=4) and with an average delay of 35.5 seconds (± 7.24 S.E.M; n=4). Following P2X7 antagonist administration, hypothalamic recordings showed a voltage deflection with an amplitude of 1.33 mV (± 0.78 S.E.M; n=4) and with a latency of 100.5 sec (± 18.1 S.E.M; n=4). P2X7 antagonism significantly increased the latency of hypothalamic voltage deflection following CSD (p=0.01) (Figure 2d-e).

### 4) CSD results in an increased neuronal activity in hypothalamus

We assessed hypothalamic activation following CSD, by counting hypothalamic c-fos positive neurons. Following CSD there was a significant increase in total number of c-fos positive neurons bilaterally in the hypothalamus (p = 0.04). This increase in c-fos positivity was prevented by P2X7R antagonist administration (p = 0.04) (Figure 2f-g). In none of the groups, c-fos positive cell number differs between the two hemispheres (the hemisphere where CSD was induced and the contralateral one) (p > 0.99; n=3/group).

### 5) P2X7 antagonism halts neuroinflammation following CSD

Following CSD, in the cortical and the subcortical structures (striatum, hippocampus, thalamus and hypothalamus), we observed an increase in the nuclear translocation of NFκB -p65 in S100ß positive astrocytes and a decrease in neuronal HMGB1 positivity (HMGB1 release) supporting previous studies^5^ (p < 0.001; n=3/group) (Figure 3a-g, Figure 4a-g; Supplemental figure 1a-c, Supplemental figure 2a-c). P2X7 antagonism inhibited the translocation of NFκB-p65 to the nucleus of astrocytes and prevented HMGB1 release from neurons significantly thus inhibiting the neuroinflammation in both hemispheres (p < 0.001, p < 0.05; n=3/group) (Figure 3c-g; Figure 4c-g; Supplemental figure 1a-c; Supplemental figure 2a-c). There was no difference in NFκB-p65 nuclear translocation or in HMGB1 release between the two hemispheres (p = 0.98).

**Figure 3.**
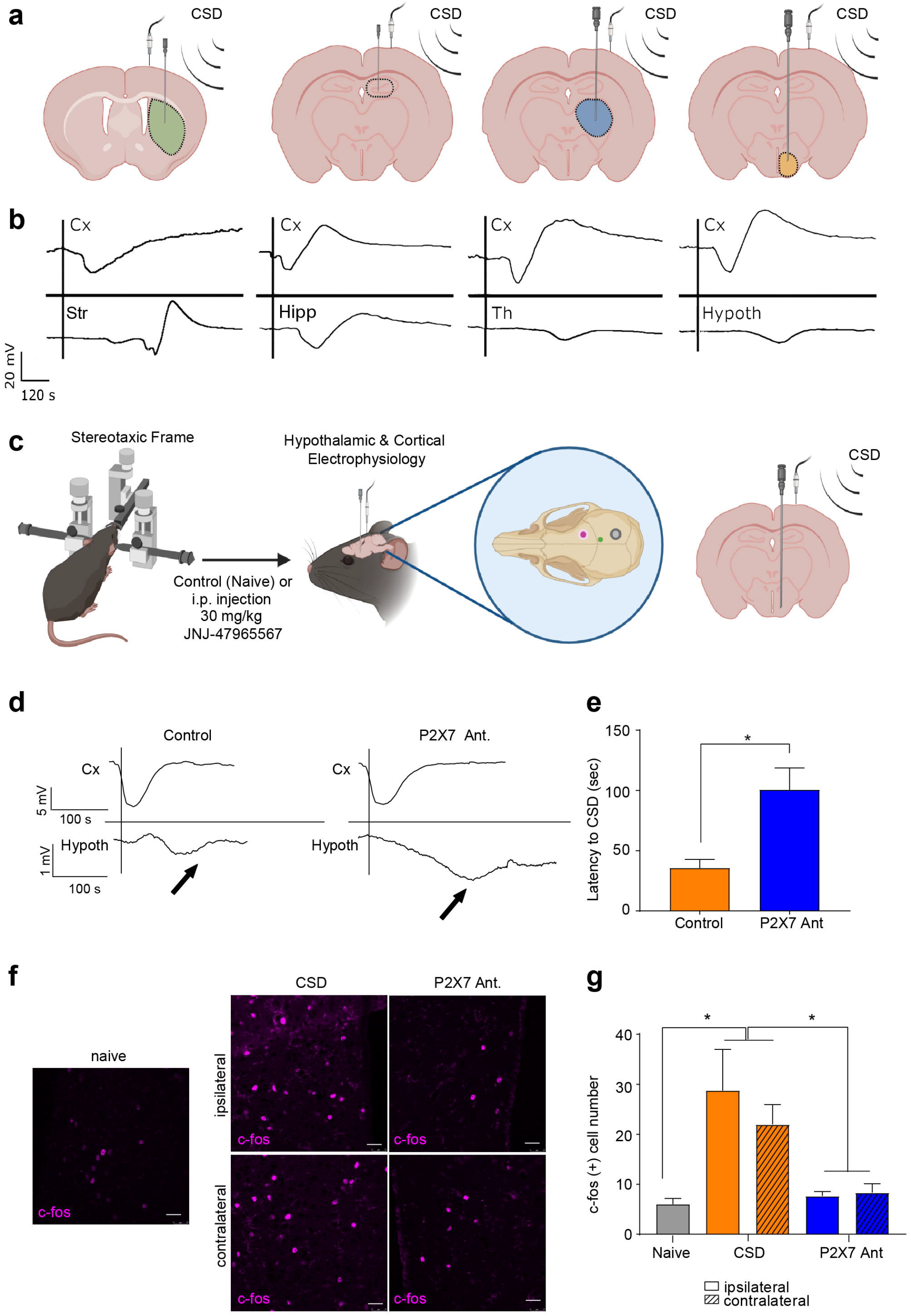
P2X7 antagonism prevents nuclear translocation of NFκB-p65 in astrocytes following CSD. **a.** Representative images of cortical NFκB-p65 and S100β immunofluorescent co-staining in naïve, sham mice, and in mice following optogenetic CSD induction with or without P2X7 antagonist administration. *scale bar:25 μm* **b.** Representative images of hypothalamic NFκB-p65 and S100ß immunofluorescent co-staining. *scale bar:25 μm* **c.** Percentage of nuclear translocation of NFκB-p65 in S100β-positive astrocytes in cortex in naïve, sham mice, and in mice following optogenetic CSD induction with or without P2X7 antagonist administration (n=3/group). **d.** Percentage of nuclear translocation of NFκB-p65 in S100β-positive astrocytes in hypothalamus (n=3/group). **e.** Percentage of nuclear translocation of NFκB-p65 in S100β-positive astrocytes in thalamus (n=3/group). **f.** Percentage of nuclear translocation of NFκB-p65 in S100β-positive astrocytes in hippocampus (n=3/group). **g.** Percentage of nuclear translocation of NFκB-p65 in S100β-positive astrocytes in striatum (n=3/group). *CSD: Cortical Spreading Depression (ns: p>0.05, *:p003C;.05, *:p003C;.01, *:p<0.001)*

**Figure 4.**
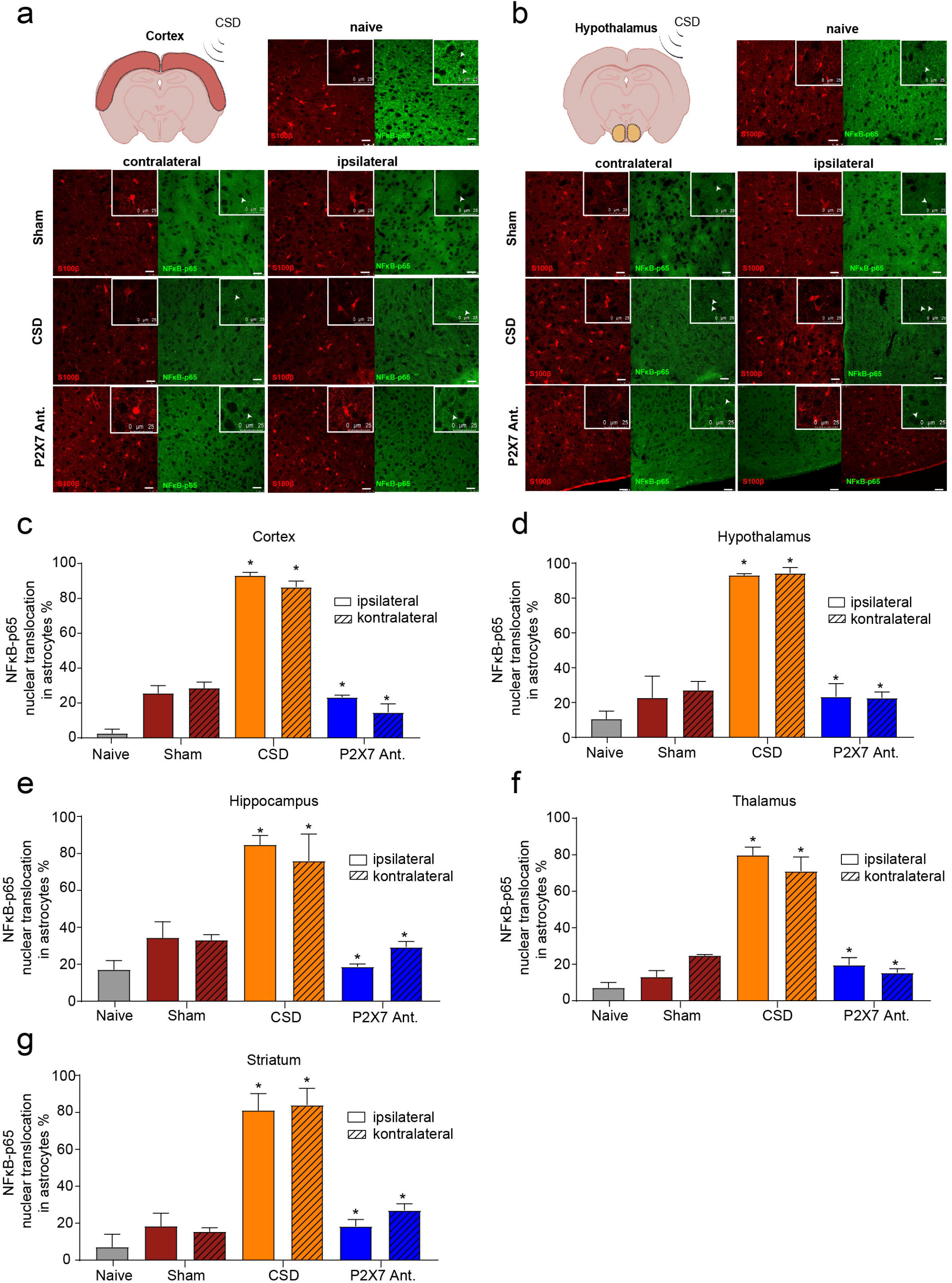
P2X7 antagonism prevents HMGB1 release following CSD. **a.** Representative images of cortical HMGB1 immunofluorescent staining in naïve, sham mice, and in mice following optogenetic CSD induction with or without P2X7 antagonist administration. *scale bar:25 μm* **b.** Representative images of hypothalamic HMGB1 immunofluorescent staining. *scale bar:25 μm* **c.** Percentage of cortical HMGB1 release in naïve, sham mice, and in mice following optogenetic CSD induction with or without P2X7 antagonist administration (n=3/group). **d.** Percentage of hypothalamic HMGB1 release (n=3/group). **e.** Percentage of thalamic HMGB1 release(n=3/group). **f.** Percentage of striatal HMGB1 release (n=3/group). **g.** Percentage of hippocampal HMGB1 release (n=3/group). *CSD: Cortical Spreading Depression (ns: p>0.05, *:p<0.05, *:p<.01, *:p<0.001)*

### 6) P2X7 receptor signal is increased in both cortical and subcortical structures following CSD and is colocalized to neurons

In the naïve mice, we observed a membranous labeling of P2X7R whereas following CSD, there was a substantial increase in cytoplasmic signal. This increase in P2X7R signal was prominent in both hemispheres. This increase in the P2X7 signal fluorescence intensity was significant in cortex, striatum, thalamus, hypothalamus and hippocampus (p < 0.001, p = 0.02, p = 0.04, p = 0.004, p = 0.0003, respectively; n=3/group) (Figure 5a-g; Supplemental figure 3a-c). Between the two hemispheres in the subcortical structures the increase in P2X7R signal was not different however in the cortex, this increase was more substantial in the ipsilateral hemisphere than the contralateral one (p = 0.004). P2X7 antagonist application prevented this P2X7 signal intensity increase following CSD, in all of the investigated brain regions, cortex (p < 0.001), striatum (p = 0.003), thalamus (p = 0.009), hypothalamus (p=0.004) and hippocampus (p = 0.004). Following P2X7 administration, the decrease in P2X7 signal intensity was bilateral in both hemispheres (Figure 5a-g; Supplemental figure 3a-c).

**Figure 5.**
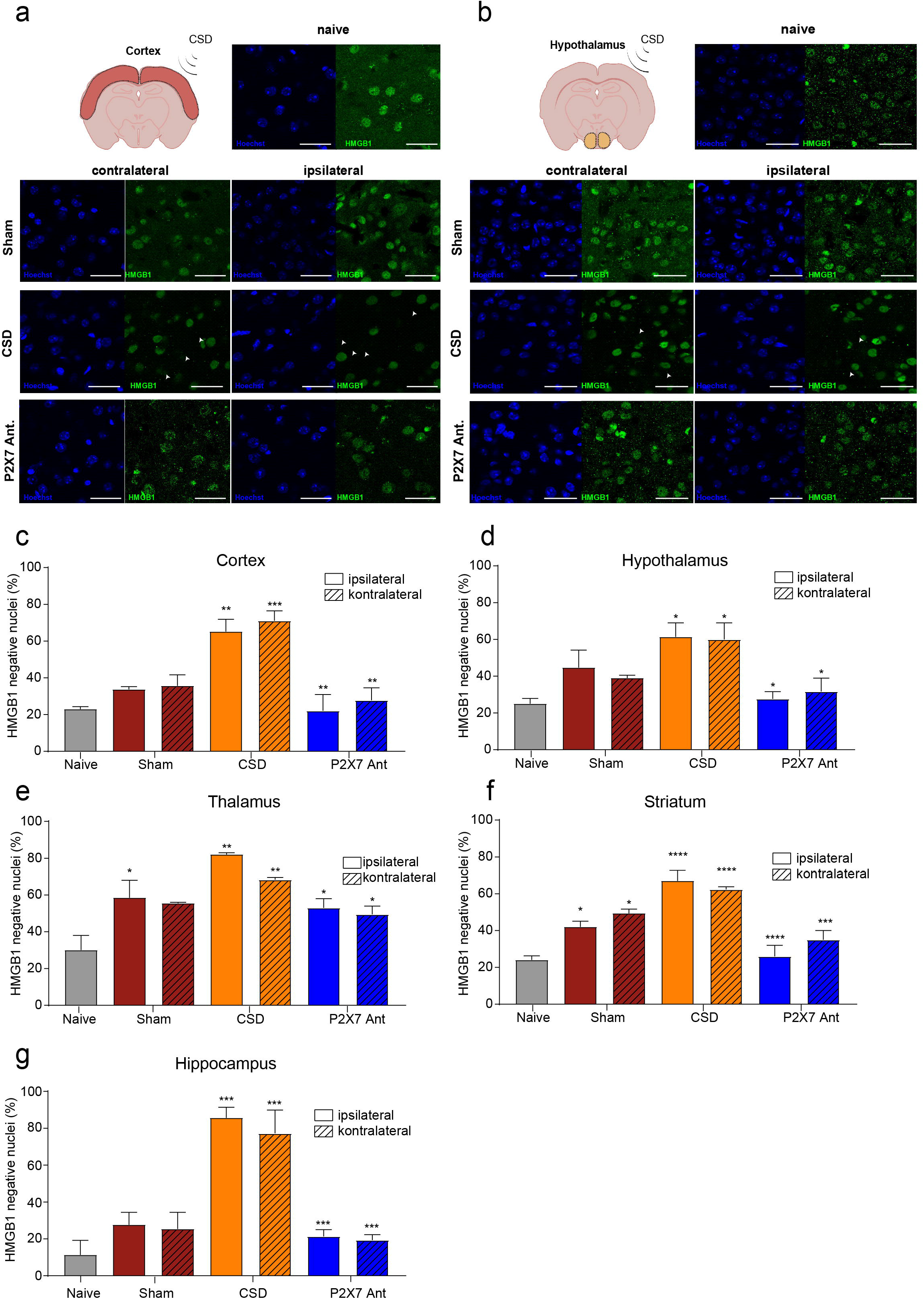
P2X7R signal is increased in cortical and subcortical structures that localizes to neurons and is prevented by P2X7 antagonism. **a.** Representative images of cortical P2X7R immunofluorescent staining in naïve mice and in mice following optogenetic CSD induction with or without P2X7 antagonist administration. *scale bar:25 μm* **b.** Representative images of hypothalamic P2X7R immunofluorescent staining. *scale bar:25 μm* **c.** Fluorescence intensity of cortical P2X7R signal in naïve mice and in mice following optogenetic CSD induction with or without P2X7 antagonist administration (n=3/group). **d.** Fluorescence intensity of hypothalamic P2X7R signal (n=3/group). **e.** Fluorescence intensity of striatal P2X7R signal (n=3/group). **f.** Fluorescence intensity of thalamic P2X7R signal (n=3/group). **g.** Fluorescence intensity of hippocampal P2X7R signal (n=3/group). **h.** Representative images of P2X7R signal following CSD co-stained with either NeuN or S100β. **i.** Pearson coefficient of P2X7R-S100β or P2X7R-NeuN colocalization (n=3/group). *(ns: p>0.05, *:p<0.05, *:p<0.01, *:p<0.001)*

We co-stained these sections with P2X7 and either NeuN (neuron marker) or S100β (astrocyte marker) to determine the cell type in which P2X7 signal increase takes place (Figure 5h). Cytoplasmic P2X7 signal colocalized to NeuN significantly (p=0.03; n=3/group) (Figure 5i).

## Discussion

Migraine is an episodic headache disorder and its pathophysiology is yet to be fully elucidated^1^. CSD is the electrophysiological equivalent of migraine aura which results in opening of Panx-1 megachannels, neuronal HMGB1 release and induction of neuroinflammatory cascades in astrocytes that eventually result in neurogenic inflammation, trigeminal activation and headache^5^. CSD was long studied as a cortical phenomenon. First time in 1964, Fifkova et al. recorded the striatal spread of the CSD using electrophysiology^11^. During a migraine attack, many symptoms could be explained by the subcortical spread of the CSD or by possible de novo spreading depressions in these subcortical structures^9^. Among these symptoms, alterations in the consciousness can be explained by thalamic spreading depression; alterations in the locomotion can be explained by striatal spreading depression; dysphoria, yawning and fluid retention can be consequences of hypothalamic spreading depression^9,17,18^ The autonomic symptoms which are more pronounced in the prodromal phase of a migraine attack, could be an effect of subcortical spreading depressions causing hypothalamic dysfunction^19^. Previously, the subcortical spread of the CSD was investigated using two different strains of Familial Hemiplegic Migraine 1 (FHM1) transgenic mice. These mice have mutations in their CACNA1A gene resulting in mutant P/Q-type voltage gated calcium channels, R192Q or S218L missense mutations, respectively. These two mutants demonstrate different levels of cortical hyperexcitability and phenotypical severity (more severe in S218L mice), and authors found that the level of subcortical spread in these mice was correlated with the phenotypical severity of their mutation^9^. A further MRI study using S218L FHM1 mice confirmed that the CSD spreads from cortex to striatum and from hippocampus to thalamus^10^. The difference in the level of subcortical spread between these two different transgenic strains (R192Q and S218L) is explained by the differential increase in cortical hyperexcitability which overcomes the barriers such as low neuronal densities or white matter, that potentially halt CSD’s spread^9^. In S218L FHM1 mice, subcortical spread of CSD could be prevented by intraperitoneal guanosine application^9^, which stimulates astrocytic glutamate reuptake, providing further evidence that extracellular glutamate, K^+^ and autacoid factors aid in the subcortical spread of CSD. In our study, we recorded waves of spreading depression in different subcortical brain areas including striatum, hippocampus, thalamus and hypothalamus after a cortical pinprick-induced CSD. This phenomenon has been noticed since early in vivo studies, however, the reported degree of penetration was variable and depended on the anesthetic that is used, in addition to other experimental parameters such as the mode of CSD induction and the method of detection ^8,20–22^. We observed an extensive subcortical spread when urethane, an anesthetic that does not significantly suppress cortical excitability, was used. Simultaneous electrophysiological recordings showed that the CSD spreads to subcortical areas with a considerable delay, supporting the original idea that it spreads via grey matter continuity ^8,23^ rather than axonal conduction, which is much faster. These findings further raise the question if there are other extracellular factors that aid the spread of CSD to the subcortical structures.

CSD opens neuronal Panx-1 megachannels and causes an increase in extracellular ATP concentration^5,7^. Released ATP can potentially be an autacoid factor that aids in the subcortical spread of the CSD as evidence supports functional coupling of Panx-1 and P2X7 receptors ^7,24^. Here we used a potent and selective, BBB permeable P2X7 antagonist, JNJ-47965567, to investigate the effects of P2X7 antagonism on CSD’s subcortical spread^15^. P2X7 antagonist administration didn’t change CSD characteristics or CSD threshold in line with the literature^7^. Chen et al. showed that selective pharmacological inhibition of the P2X7 channel did not affect spreading depolarization threshold or frequency^7^. However, systemic or topical administration of A-438079 and BBG which block P2X7/Panx1 complex, was found to increase the CSD threshold^7^. This suggests that Panx-1 megachannels might serve as an important target in the induction and the spread of the CSD as the crucial step before P2X7 receptor activation^5^. It is also well known that purinergic receptors form heteromeric complexes and besides P2X7 receptor there are other purinergic receptors including P2X2, P2X4 that could response to extracellular ATP^25–29^. The reason why P2X7 antagonism did not have an effect on the CSD threshold might be due to the compensatory mechanisms elicited by other purinergic receptors in response to an increase in extracellular ATP. P2X7 antagonism, however, increased the latency of hypothalamic voltage deflection following CSD. This voltage deflection that we observe at hypothalamic recordings could also be a reflection of the spread of the CSD to amygdala or to the piriformis cortex on the electrode. Regardless, the significant increase in the latency suggests that extracellular ATP acts as a mediator in the subcortical spread of CSD. This finding supports previous studies that show modulation of neuronal excitability through P2X7 activation^30,31^.

In the literature, CSD is conventionally induced by electrical stimulation, topical KCl application or pin-prick, which have non-specific glial and vascular effects^14^. Following the utilization of optogenetics, CSD was successfully induced without disrupting integrity of the skull^14^. This method is particularly useful to study inflammatory pathways associated with CSD, discarding the non-specific inflammation associated with craniotomy. Here, we used Thy1-ChR2 transgenic mice and stimulated the motor cortex via a blue laser (450 nm) to induce CSD. The optogenetic CSD threshold along with other CSD characteristics (amplitude and normalized cumulative change in voltage) were determined and didn’t differ between homozygous and heterozygous transgenic animals which is in line with the literature^32^.

In various studies, c-fos (an early neuronal activation marker) is used as an indicator of CSD spread to the subcortical structures^9^. In the wild-type mice and R192Q and S218L-mutant FHM1 mice there was an increase in cortical c-fos positivity and in S218L mice there was an increase in c-fos positive neurons unilaterally in hippocampus, bilaterally in thalamic and lateral hypothalamic nuclei^9^. In another study, following CSD induction in freely-moving rats, abundant c-fos positivity was observed in the thalamic reticular nuclei and magnocellular area of hypothalamus^33^. Same study also showed that anesthesia application (thiopental, chloral hydrate) decreased aforementioned c-fos positivity, further suggesting that anesthesia alone can halt subcortical spread of CSD^33^. In our study, we found a bilateral increase in the c-fos positive neuron number in hypothalamus following CSD. P2X7 administration however, prevented this increase in the c-fos positive neuron number, pointing out the role of P2X7 receptors in the hypothalamic neuronal activation following CSD.

CSD opens neuronal Panx-1 megachannels and induces neuroinflammation^5^. Opening of Panx-1 channels results in an increase in extracellular ATP concentration^7^. Panx-1 channels are also known to be in close relation to the P2X7 receptors on the membrane and that P2X7 receptor activation triggers the inflammatory pathways rendering P2X7 as a potential target in neuroinflammation^34,35^. In our study, P2X7 antagonism prevented the nuclear translocation of NFκB-p65 and HMGB1 release in cortex, thalamus, striatum, hippocampus and hypothalamus following optogenetically induced CSD. This finding indicates that the opening of the Panx-1 channels following CSD induces inflammation via a P2X7 receptor-mediated mechanism triggered by increased extracellular ATP. In another study, supporting our finding, application of P2X7/Panx1 complex blockers, A-438079 and BBG preceded by CSD was shown to decrease cortical IL 1β mRNA levels^7^. Interestingly, we didn’t only see the effects of P2X7 antagonism on neuroinflammation in the ipsilateral hemisphere where CSD was induced but also in the contralateral hemisphere. A recent clinic PET-MRI study confirmed this finding which was conducted with patients that had at least 1 migraine attack with aura and healthy controls, used a (^11^C)PBR28 radio ligand that binds to a glial marker 18 kDa translocator protein which increases during neuroinflammation, showed an increase in inflammatory signal in cortical and subcortical structures bilaterally in migraine patients and the signal intensity was found to correlate with the number of migraine attacks^36^.

P2X7 receptors play an important role in the pathophysiology of various neuropsychiatric diseases and it has been long debated whether neuronal P2X7 receptors existed, if neuronal or glial P2X7R play a role in certain neurological and psychiatric disease pathophysiologies^37,38^. In 2017, Mancarci et al presented an integrated mouse single-cell RNA transcriptomics database, which proved the neuronal presence of P2X7 receptors^39^. Further studies confirmed the neuronal presence of P2X7 and showed that the roles of these receptors in the glia or neurons are not always mutually exclusive^40^. Here, we investigated P2X7 immunostaining signal following CSD and we have found an increase in the P2X7 signal in cortex, thalamus, striatum, hypothalamus and hippocampus following CSD in neurons, effecting both hemispheres, concordant with the neuroinflammation^41^. In addition, P2X7 antagonism thus blockage of the inflammatory cascade halted this increase in the neuronal cytoplasmic P2X7 signal in all of the aforementioned cortical and subcortical regions. Previous studies show that P2X receptors are internalized following increased extracellular ATP, as an adaptive response to protect the neurons from ATP-related toxicity ^42^. This protective mechanism exists in various agonist-receptor combinations when high extracellular concentrations of the agonist has toxic effects to the cell^42,43^. Acute internalization of the P2X7 receptors, 20 minutes following the CSD, explains the increased cytoplasmic P2X7R signal. Meanwhile, P2X7 antagonist administration prevents this adaptive response, supporting our findings explaining the concordance between neuroinflammation and P2X7 signal increase following CSD^44^.

## Conclusion

In conclusion, this study shows that P2X7 receptors have an important role in neuroinflammation in cortical and subcortical structures (striatum, thalamus, hippocampus, hypothalamus) and in hypothalamic neuronal activation in both hemispheres following CSD. Moreover, P2X7R antagonism increased the latency of hypothalamic voltage deflection following CSD, pointing out the role of extracellular ATP as a mediator in the subcortical spread of CSD. CSD also caused an increase in the P2X7 signal that may be secondary to receptor internalization as an adaptive response to protect neurons from ATP-associated toxicity. These results suggest that P2X7 receptors can be used as a potential target in migraine treatment or other neurological diseases where CSD takes place.

## Methods

### 1) Animals

In this study, C57BL6 male, wild-type and/or transgenic mice (Thy1-ChR2) weighting 20-25 grams were used. Standard housing conditions were applied, in 12 hours light/dark cycles, at 22°C room temperature and ad libitum. All the experiments performed on laboratory animals are approved by Hacettepe University Animal Studies Ethical Committee (2008/55-6, 2010/49-1 and 2017/11-5).

### 2) Electrophysiology

#### 2.1 Surgery

Before the surgical procedures, mice were anesthesized with xylazine (10 mg/kg, Alfazyne, 2% Alfasan, Netherlands) and Urethane (1.25 g/kg, Sigma-Aldrich, USA). Depth of the anesthesia was tested with a pinch to the paw and animals were transferred to the stereotaxic frame. During the experiments, animals were supplied with 2L/min oxygen and followed up with spontaneous respiration. A rectal probe was used during the procedure and the body temperature was kept at 37±2°C with a homeothermic blanket. Meanwhile blood oxygen saturation was monitorized with a pulse oximetry.

After placing the mice in the stereotaxic frame, a skin incision was made under a stereomicroscope to expose bregma, parietal and frontal bones. A region on the posterior parietal bone was thinned using a drill for electrode placement. During drilling, cold saline solution was used to cool down the skull. Optogenetic laser stimulation was performed transcranially on the frontal bone (on motor cortex) while electrophysiological recording was made with an Ag-AgCl covered 1 mm pellet electrode on the posterior parietal bone. In the sham group the same surgical procedure was performed with wild-type mice, where CSD was never induced with laser stimulation.

#### 2.2 Optogenetic CSD Threshold

Following the surgical procedures, the fiberoptic cable was placed on the frontal bone and optogenetic threshold experimental protocol was performed according to the method described in (Houben et al). Laser source was adjusted to 4 mW using a photometer and 450 nm laser was applied step by step for 0.25; 0.5; 0.75; 1; 2; 3; 4; 5; 6; 7; 8; 9; 10; 11; 12; 13; 14; 15 seconds. After each stimulation, there was a wait of 5 minutes before the next stimulation to see if CSD is triggered where extracranial electrophysiological recordings were performed simulatenously. The energy transferred to the cortex was calculated as milijoules (mJ= mW x sec). After induction of CSD the experiment was terminated after 20 minutes.

#### 2.3 Subcortical Electrophysiological Recordings

So as to investigate the subcortical spread of the CSD, following opening a burr hole at the coordinates according to the bregma a tungsten electrode with a 1 μm tip diameter was placed to dentate gyrus (2 mm posterior, 1.5 mm lateral to bregma, 1.8 mm deep from the surface), to thalamus (ventral posterolateral nucleus, 1.6 mm posterior, 1.75 mm lateral to bregma, 3.25 mm in deep), to striatum (0.86 mm anterior, 1.5 mm lateral to bregma, 3 mm in deep), and to hypothalamus (anterior hypothalamic area, 1.46 mm posterior, 0.5 mm lateral, 5.5 mm in deep from the surface). After the placement of the electrode, there was a wait of 20 minutes before inducing CSD by pin-prick using a frontal burr hole. Electrophysiological recordings from a surface electrode was performed simultaneously. These experiments were performed primarly by the pin-pick method since subcortical electrophysiological recordings are invasive although they were replicated via optogenetic induction of CSD. The pin-prick method was not used in investigating the neuroinflammation following CSD. P2X7R antagonist or vehicle was administered before CSD induction during simultaneous hypothalamic recordings.

### 3) Pharmacological Agent

A highly potent (pK_i_:7.9±0.07), selective (EC_50_:78±19 ng.ml^-1^) and BBB permeable P2X7 antagonist, JNJ-47965567 was used to investigate the effects of P2X7 antagonism on the subcortical spread of CSD and neuroinflammation. The pharmacokinetic properties of this drug was characterized in the literature indicating that the drug concentration peaks rapidly in the brain (in 15 minutes) and stays effective for 4-6 hours ^45^. Drug was freshly prepared for each experiment and dissolved in 30% solution of cyclodextrin-sulfobutyl ether sodium salt, administered intraperitoneally 15 minutes before the CSD threshold protocol started.

### 4) Immunostaining methods

Following the in vivo experiments, all mice went under cardiac perfusion with 0.4% heparin followed by 4% paraformaldehyde (PFA). The brains were extracted and incubated in 4% PFA for 24 h then cryoprotected with 30% sucrose. Coronal sections of 20 μm were made using a cryostat and these sections were further used for immunostaining. Two different methods of immunostaining were used: indirect immunofluorescence, immunosignal hybridization chain reaction (isCHR)

#### 4.1. Indirect Immunofluorescence

First, antigen retrieval was performed at 80°C for 10 minutes using citrate buffer (pH 6.0 and sections were washed with phosphate buffered saline (PBS) prior to blockage. Sections were blocked in room temperature with 10% normal goat serum (NGS)-0.5% Triton X-0.3 M glycine for an hour at room temperature. Then the sections were incubated with primary antibody of interest in the blockage solution (c-fos,1:200, Abcam/ab208942; NFκB-p65, 1:200, Cell Signaling Technology/8242; P2X7R, 1:50, Abcam/ab109054; S100β, 1:200, Abcam/ab52642; NeuN, 1:200, Millipore/MAB377) overnight at +4°C. The day after sections were washed 3 times with PBS and incubated with secondary antibodies (Goat anti-rabbit Cy2, 1:200, Jackson Immunoresearch (JI)/111-225-144; Goat anti-rabbit Cy3, 1:200, JI /111-165-144; Goat anti-mouse Cy2, 1:200, JI /115-225-146; Goat anti-mouse Cy3, 1:200, JI /115-165-146) in blockage solution for 2 hours in room temperature. After washing the sections with PBS three times, the sections were mounted with Hoechst-33258 and imaged under confocal microscope

#### 4.2 Immunosignal Hybridization Chain Reaction (isHCR)

isHCR is a technique developed from fluorescent in situ hybridization (FISH), that amplifies the fluorescent signal using complementary primers which have fluorescent tags. This system was used with NFκB-p65 staining where spatial information is vital and signal is rather difficult to obtain. Following the incubation with the primary antibody the section was incubated with a secondary antibody that has a biotin tag. Using the strong affinity of streptavidin and biotin, streptavidin was used as a linker between the secondary antibody and the initiator primer sequence tagged with biotin. Then the amplifier sequences which have a fluorescent tag were used to amplify the signal. Background signal was reduced using graphene oxide. The details of the method is reviewed in^46^.

### 5) Analysis of the Confocal Images

Following NFκB-p65 and P2X7 staining, 2 images per hemisphere were taken for each brain region (cortex, striatum, hippocampus, hypothalamus, thalamus) per each animal. For analysis of the NFκB-p65 staining, total number of S100β positive cells (astrocytes) and the ones where NFκB-p65 was translocated to the nucleus were counted. For analysis of the P2X7R staining, cellular fluorescent intensity was investigated via ImageJ after the background was subtracted from each image. Then the colocalization of P2X7 with S100β or NeuN was made so as to determine the cellular subtype where the increase in the signal was observed. Following c-fos staining 3 images were taken from hypothalamus from each side and the c-fos positive cells were counted.

### 6) Statistical Analysis

For the statistical analysis IBM SPSS 20 (Statistical Package for Social Sciences) and Graphpad Prism were used. When more than two independent group were compared ANOVA test or Kruskal-Wallis test was used decided upon the normal distribution parameters of the data. If the result was significant, appropriate post-hoc tests were used with Bonferroni correction. When normal distribution parameters were not met, Mann Whitney U test was used.

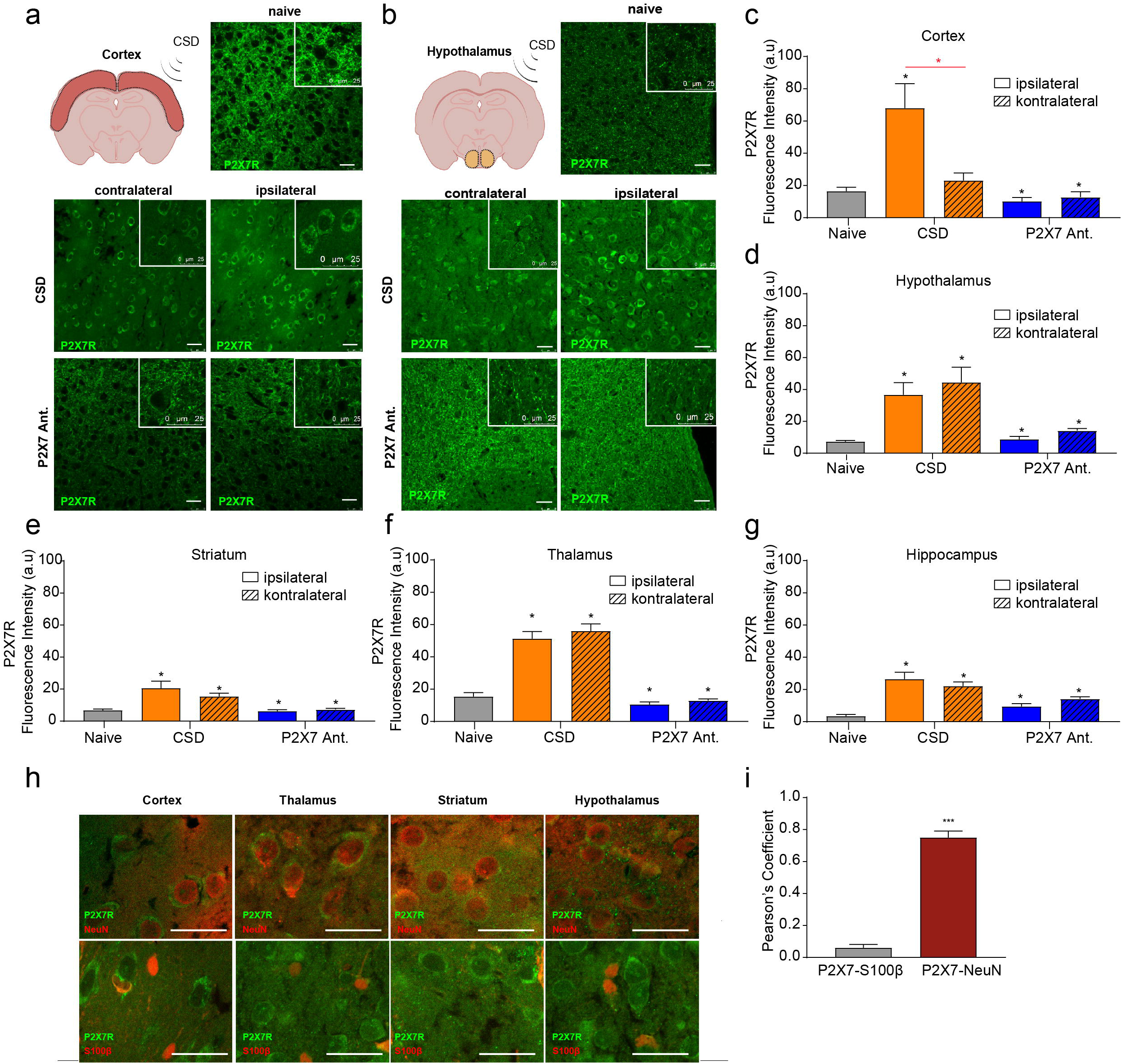

## References

1. Lipton RB, Diamond S, Reed M, et al. Migraine diagnosis and treatment: Results from the American migraine study II. Headache 2001;41(7):638–645.

2. Leão AAP, Morison RS. PROPAGATION OF SPREADING CORTICAL DEPRESSION. [Internet]. J. Nerv. Ment. Dis. 1945;102(5) Available from: http://journals.lww.com/jonmd/Fulltext/1945/11000/PROPAGATION_OF_SPREADING_CORTICAL_DEPRESSION_.29.aspx

3. Olesen J, Burstein R, Ashina M, Tfelt-Hansen P. Origin of pain in migraine: evidence for peripheral sensitisation [Internet]. Lancet Neurol. 2017;8(7):679–690. Available from: http://dx.doi.org/10.1016/S1474-4422(09)70090-0

4. Hadjikhani N, Sanchez Del Rio M, Wu O, et al. Mechanisms of migraine aura revealed by functional MRI in human visual cortex. Proc. Natl. Acad. Sci. U. S. A. 2001;98(8):4687–4692.

5. Karatas H, Erdener SE, Gursoy-Ozdemir Y, et al. Spreading depression triggers headache by activating neuronal Panx1 channels. Science 2013;339(6123):1092–1095.

6. Lister MF, Sharkey J, Sawatzky DA, et al. The role of the purinergic P2X7 receptor in inflammation. J. Inflamm. (Lond). 2007;4:5.

7. Chen S-P, Qin T, Seidel J, et al. Inhibition of the P2X7-PANX1 complex suppresses spreading depolarization and neuroinflammation. Brain 2017;140

8. Fifkova E, Syka J. Relationships between cortical and striatal spreading depression in rat [Internet]. Exp. Neurol. 1964;9(5):355–366. Available from: http://linkinghub.elsevier.com/retrieve/pii/0014488664900706

9. Eikermann-Haerter K, Yuzawa I, Qin T, et al. Enhanced subcortical spreading depression in familial hemiplegic migraine type 1 mutant mice. J. Neurosci. 2011;31(15):5755–5763.

10. Cain SM, Bohnet B, LeDue J, et al. In vivo imaging reveals that pregabalin inhibits cortical spreading depression and propagation to subcortical brain structures. Proc. Natl. Acad. Sci. U. S. A. 2017;114(9):2401–2406.

11. Maniyar FH, Sprenger T, Monteith T, Schankin C. Brain activations in the premonitory phase of nitroglycerin-triggered migraine attacks. Brain 2014;137(2013):232–241.

12. Goadsby PJ, Edvinsson L, Ekman R. Vasoactive peptide release in the extracerebral circulation of humans during migraine headache. Ann. Neurol. 1990;28(2):183–187.

13. Megirian D. Unilateral cortical spreading depression and conditioned eye blink responses in rabbits. Acta Neurobiol. Exp. (Wars). 1973;33(4):699–710.

14. Houben T, Loonen IC, Baca SM, et al. Optogenetic induction of cortical spreading depression in anesthetized and freely behaving mice. [Internet]. J. Cereb. Blood Flow Metab. 2016;Available from: http://www.ncbi.nlm.nih.gov/pubmed/27107026

15. Bhattacharya A, Wang Q, Ao H, et al. Pharmacological characterization of a novel centrally permeable P2X7 receptor antagonist: JNJ-47965567. Br. J. Pharmacol. 2013;170(3):624–640.

16. Merkler D, Klinker F, Jürgens T, et al. Propagation of spreading depression inversely correlates with cortical myelin content. Ann. Neurol. 2009;66(3):355–365.

17. Cutrer FM OJ. Migraines with aura and their subforms. In: The headaches, Ed 3. Philadelphia: Lippincott Williams and Wilkins.; 2006 p. pp 407–422.

18. Fu X, Chen M, Lu J, Li P. Cortical spreading depression induces propagating activation of the thalamus ventral posteromedial nucleus in awake mice. 2022;1–10.

19. Mraovitch S, Calando Y, Goadsby PJ, Seylaz J. Subcortical cerebral blood flow and metabolic changes elicited by cortical spreading depression in rat. Cephalalgia 1992;12(3):137–141.

20. Křivánek J, Fifková E. The value of ultramicro-analysis of lactic acid in tracing the penetration of Leão’s cortical spreading depression to subcortical areas [Internet]. J. Neurol. Sci. 1965;2(4):385–392. Available from: https://www.sciencedirect.com/science/article/pii/0022510X65900201

21. Arabia A-M, Shen P-J, Gundlach AL. Increased striatal proenkephalin mRNA subsequent to production of spreading depression in rat cerebral cortex: activation of corticostriatal pathways? [Internet]. Mol. Brain Res. 1998;61(1):195–202. Available from: https://www.sciencedirect.com/science/article/pii/S0169328X98001892

22. Eikermann-Haerter K, Dileköz E, Kudo C, et al. Genetic and hormonal factors modulate spreading depression and transient hemiparesis in mouse models of familial hemiplegic migraine type 1 [Internet]. J. Clin. Invest. 2009;119(1):99–109. Available from: https://doi.org/10.1172/JCI36059

23. Křivánek J, Fifková E. The value of ultramicro-analysis of lactic acid in tracing the penetration of Leão’s cortical spreading depression to subcortical areas. J. Neurol. Sci. 1965;2(4):385–392.

24. Bravo D, Zepeda-Morales K, Maturana CJ, et al. NMDA and P2X7 Receptors Require Pannexin 1 Activation to Initiate and Maintain Nociceptive Signaling in the Spinal Cord of Neuropathic Rats [Internet]. Int. J. Mol. Sci. 2022;23(12) Available from: https://www.mdpi.com/1422-0067/23/12/6705

25. Roberts JA, Vial C, Digby HR, et al. Molecular properties of P2X receptors. Pflugers Arch. 2006;452(5):486–500.

26. Guo C, Masin M, Qureshi OS, Murrell-Lagnado RD. Evidence for functional P2X4/P2X7 heteromeric receptors. Mol. Pharmacol. 2007;72(6):1447–1456.

27. Khmyz V, Maximyuk O, Teslenko V, et al. P2X3 receptor gating near normal body temperature. Pflugers Arch. 2008;456(2):339–347.

28. Abbracchio MP, Burnstock G, Boeynaems J-M, et al. International Union of Pharmacology LVIII: update on the P2Y G protein-coupled nucleotide receptors: from molecular mechanisms and pathophysiology to therapy. Pharmacol. Rev. 2006;58(3):281–341.

29. Cavaliere F, Florenzano F, Amadio S, et al. Up-regulation of P2X2, P2X4 receptor and ischemic cell death: prevention by P2 antagonists. Neuroscience 2003;120(1):85–98.

30. Ireland MF, Noakes PG, Bellingham MC. P2X7-like receptor subunits enhance excitatory synaptic transmission at central synapses by presynaptic mechanisms. Neuroscience 2004;128(2):269–280.

31. del Puerto A, Fronzaroli-Molinieres L, Perez-Alvarez MJ, et al. ATP-P2X7 Receptor Modulates Axon Initial Segment Composition and Function in Physiological Conditions and Brain Injury [Internet]. Cereb. Cortex 2014;25(8):2282–2294. Available from: https://doi.org/10.1093/cercor/bhu035

32. Chung DY, Sadeghian H, Qin T, et al. Determinants of optogenetic cortical spreading depolarizations. Cereb. Cortex 2019;29(3):1150–1161.

33. Tepe N, Filiz A, Dilekoz E, et al. The thalamic reticular nucleus is activated by cortical spreading depression in freely moving rats: prevention by acute valproate administration. Eur. J. Neurosci. 2015;41(1):120–128.

34. Munoz-Planillo R, Kuffa P, Martinez-Colon G, et al. K(+) efflux is the common trigger of NLRP3 inflammasome activation by bacterial toxins and particulate matter. Immunity 2013;38(6):1142–1153.

35. Coppi E. Purines as Transmitter Molecules: Electrophysiological Studies on Purinergic Signaling in Different Cell Systems. Premio Firenze Univ. Press Tesi Di Dottorato 2008;23(Floransa Üniversitesi, Doktora Tezi)

36. Albrecht DS, Mainero C, Ichijo E, et al. Imaging of neuroinflammation in migraine with aura: A [11 C]PBR28 PET/MRI study. Neurology 2019;92(17):e2038–e2050.

37. Diaz-Hernandez M, Diez-Zaera M, Sanchez-Nogueiro J, et al. Altered P2X7-receptor level and function in mouse models of Huntington’s disease and therapeutic efficacy of antagonist administration. FASEB J. Off. Publ. Fed. Am. Soc. Exp. Biol. 2009;23(6):1893–1906.

38. Illes P, Khan TM, Rubini P. Neuronal P2X7 Receptors Revisited: Do They Really Exist? [Internet]. J. Neurosci. 2017;37(30):7049–7062. Available from: https://pubmed.ncbi.nlm.nih.gov/28747388

39. Mancarci BO, Toker L, Pavlidis P, et al. Cross-Laboratory Analysis of Brain Cell Type Transcriptomes with Applications to Interpretation of Bulk Tissue Data. 2017;4(December)

40. Miras-Portugal MT, Sebastián-Serrano Á, de Diego García L, Díaz-Hernández M. Neuronal P2X7 Receptor: Involvement in Neuronal Physiology and Pathology [Internet]. J. Neurosci. 2017;37(30):7063 LP – 7072. Available from: http://www.jneurosci.org/content/37/30/7063.abstract

41. Dehghani A, Phisonkunkasem T, Yilmaz Ozcan S, et al. Widespread brain parenchymal HMGB1 and NF-κB neuroinflammatory responses upon cortical spreading depolarization in familial hemiplegic migraine type 1 mice. Neurobiol. Dis. 2021;156:105424.

42. Robinson LE, Murrell-lagnado RD. The trafficking and targeting of P2X receptors. 2013;7(November):1–6.

43. Ennion SJ, Evans RJ. Agonist-stimulated internalisation of the ligand-gated ion channel P2X1 in rat vas deferens [Internet]. FEBS Lett. 2001;489(2):154–158. Available from: https://www.sciencedirect.com/science/article/pii/S0014579301021020

44. Humphreys BD, Dubyak GR. Modulation of P2X7 nucleotide receptor expression by pro- and anti-inflammatory stimuli in THP-1 monocytes [Internet]. J. Leukoc. Biol. 1998;64(2):265–273. Available from: https://doi.org/10.1002/jlb.64.2.265

45. Jimenez-Pacheco A, Diaz-Hernandez M, Arribas-Blázquez M, et al. Transient P2X7 receptor antagonism produces lasting reductions in spontaneous seizures and gliosis in experimental temporal lobe epilepsy. J. Neurosci. 2016;36(22):5920–5932.

46. Lin R, Feng Q, Li P, et al. A hybridization-chain-reaction-based method for amplifying immunosignals [Internet]. Nat. Methods 2018;(September 2017) Available from: https://www.nature.com/articles/nmeth.4611

