## Supplementary Data for "The Effect of P2X7 Antagonism on Subcortical Spread of Optogenetically-Triggered Cortical Spreading Depression and Neuroinflammation"

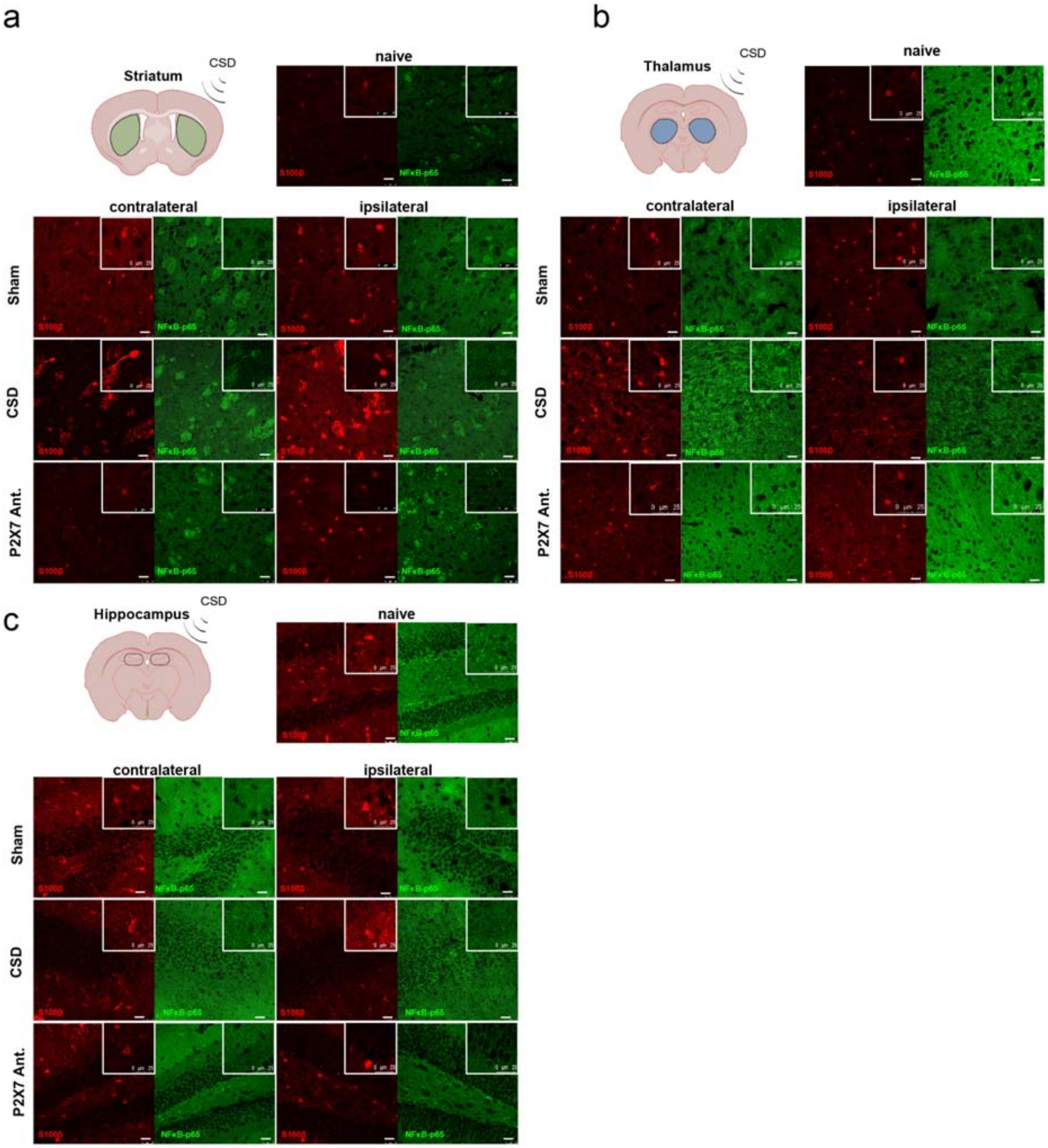


**Supplemental Figure 1 a.** Representative images of striatal NFκB-p65 and S100β immunofluorescent co-staining. *scale bar:25 μm* **b.** Representative images of thalamic NFκB-p65 and S100β immunofluorescent co-staining. *scale bar:25 μm* **c.** Representative images of hippocampal NFκB-p65 and S100β immunofluorescent co-staining. *scale bar:25 μm*

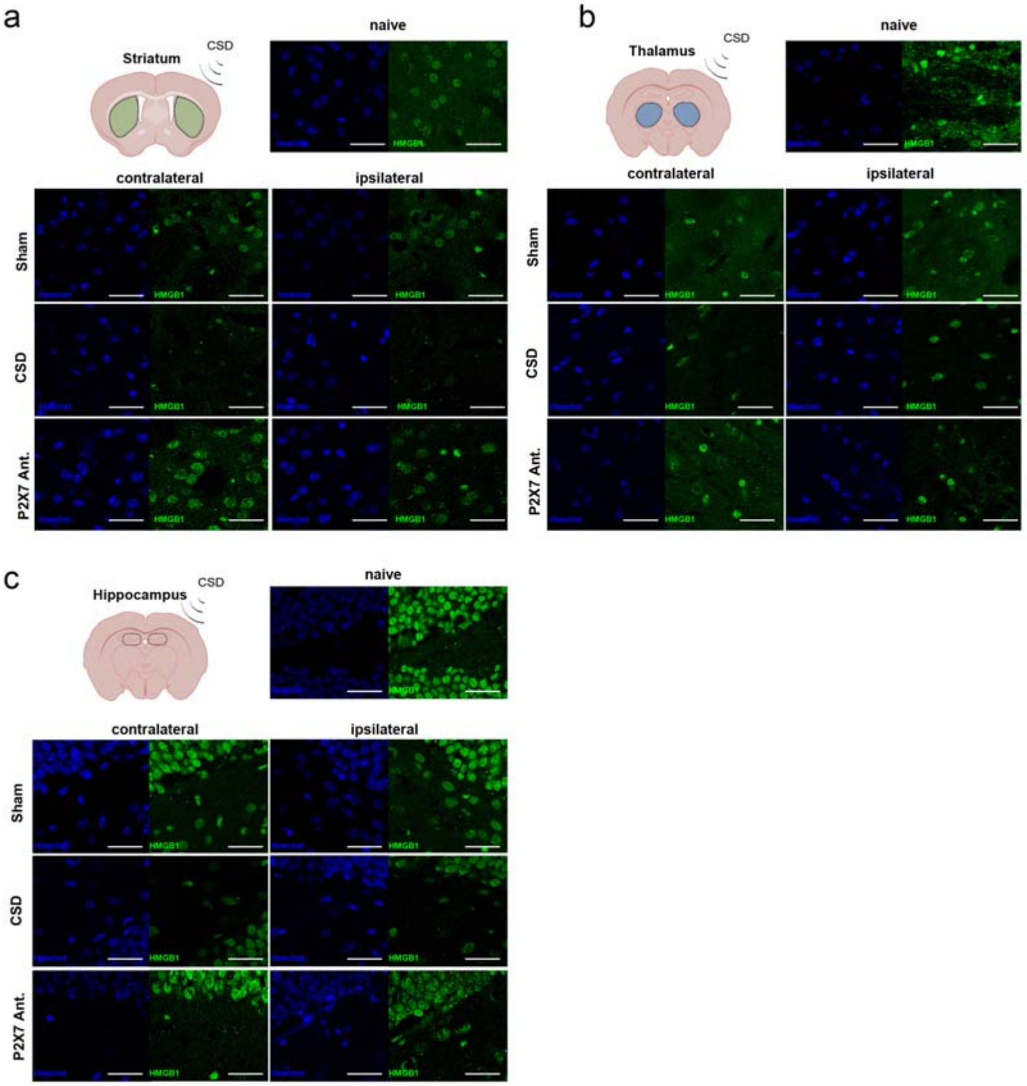


**Supplemental Figure 2 a.** Representative images of striatal HMGB1 immunofluorescent staining. *scale bar:25 $\mu m$* **b.** Representative images of thalamic HMGB1 immunofluorescent staining. *scale bar:25 $\mu m$* **c.** Representative images of hippocampal HMGB1 immunofluorescent staining. *scale bar:25 $\mu m$*

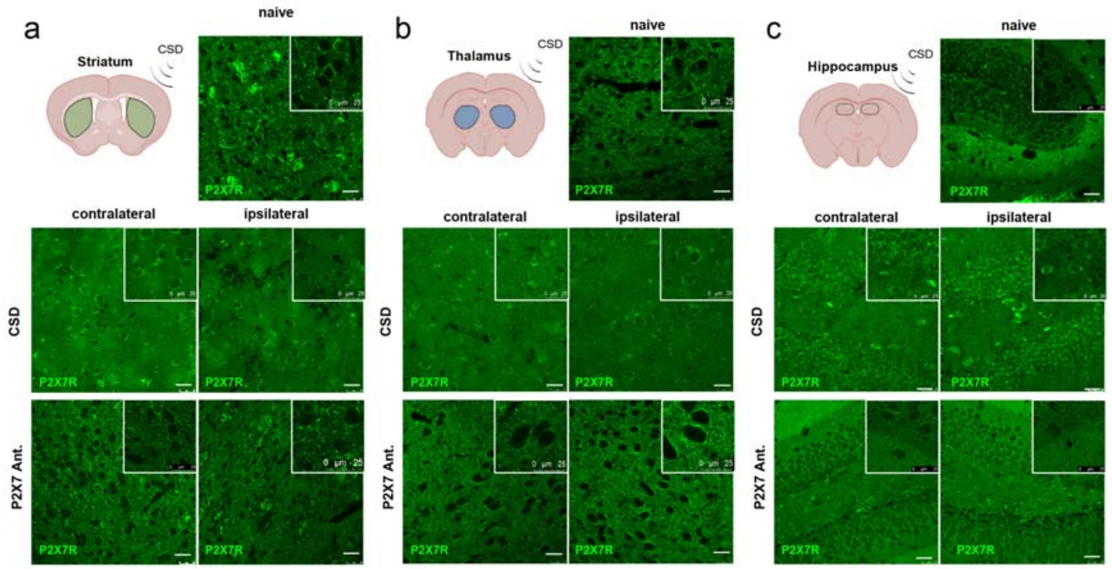


**Supplemental Figure 3 a.** Representative images of striatal P2X7R immunofluorescent staining. *scale bar:25 μm* **b.** Representative images of thalamic P2X7R immunofluorescent staining. *scale bar:25 μm* **c.** Representative images of hippocampal P2X7R immunofluorescent staining. *scale bar:25 μm*
